# Effects of two different compounds on seizure suppression using the zebrafish PTZ-seizure model

**DOI:** 10.1101/2024.07.29.605666

**Authors:** Jhonathan Angel Araujo Fernández, Thatiane Cristina de Moura, Sabela Fernández Vila, Juan Andrés Rubiolo Gaytán, Iñaki López-Díaz, Soraya Learte-Aymamí, M. Eugenio Vázquez, Maria D. Mayán, Laura Sánchez, Claudia Vianna Maurer-Morelli

## Abstract

Epilepsies are a common and severe neurological condition characterized by spontaneous and recurrent seizures. Although anti-seizure medications are effective for most patients, about 30% remain pharmacoresistant. Moreover, uncontrolled seizures are associated with risk factors and shortened life expectancy for individuals with refractory epilepsy. Preclinical studies are an essential step for drug discovery and the zebrafish (*Danio rerio*) has been successfully employed for this purpose. In this study, we applied the zebrafish PTZ-seizure model to investigate the effect of two compounds on seizure suppression, Tripeptide (p-BTX-I) and the Cx43 peptide CX2. Zebrafish larvae at 6 days post-fertilization (dpf) were exposed to both compounds, according to their group, 24h prior to PTZ-seizure induction. We quantified the compounds’ effect on seizure latency, number of seizures and transcript levels of genes related to inflammation, oxidative stress, and apoptosis (*il1b, tnfa, cox1, cox2a, il6, casp3a, casp9, baxa, bcl2a, nox1, sod1* and *cat*).

Our results showed that CX2 at a concentration of 0.1 μM/mL yielded the best outcome for seizure suppression as it reduced the number of seizures and increased the seizure latency. Additionally, CX2 treatment before PTZ-induced seizures decreased the transcript of *il1b, il6, tnfa* and *cox1* genes, all related to inflammation. A bio-distribution study showed that the CX2 reached the zebrafish brain at both times investigated, 1h and 6h. Similarly, the tripeptide exhibited anti-inflammatory and anti-apoptotic action, reducing mRNA expression of the *il1b* and *casp9* genes. Our findings suggest that both Tripeptide and CX2 hold translational potential for seizure suppression.

## Introduction

Epilepsies are a common neurological disease characterized by recurrent unprovoked seizures, affecting approximately 50 million people worldwide [1,2]. Currently, anti-seizure medications are effective for 66% of people with epilepsy in developed countries [3,4]. However, more than 30% of people with epilepsy do not respond well to conventional therapies, making them pharmacoresistant [5]. Uncontrolled seizures can lead to various risks for people with epilepsy, such as risk of injury, neuropsychological impairment, and shortened lifespan [6]. Therefore, the search for new drugs or substances that could improve the treatment of people with epilepsy is urgent.

Pre-clinical trials are crucial in testing the therapeutic or toxicological effects of new substances, using *in vivo, in vitro*, or *ex-vivo* strategies. The scalability and rapid access to the results significantly contribute to the value of such studies. In this context, the zebrafish emerges as a pivotal model, offering distinct advantages over other animal models for pre-clinical trials [7]. These include its small size, low maintenance cost, high fecundity, and optical transparency during embryogenesis [8]. Furthermore, the zebrafish genome exhibits approximately 70% homology with the human genome, with 84% of known genes associated with human diseases, including epilepsy [9,10]. In recent years, several platforms for automatic data acquisition and analysis in the zebrafish model have been developed, enabling a variety of multiplexed phenotypic assays with minimal human intervention [11,12]. Moreover, the zebrafish aligns with the 3Rs (reduction, refinement, and replacement) philosophy [13]. Beyond its investigation of molecular pathways and behaviors relevant to various diseases, the zebrafish offers a time and cost-effective model due to its small size, among other attributes, which is advantageous for scalable studies involving simultaneous analysis of animals and drugs [14].

In the context of epilepsy, the zebrafish is a well-characterized seizure model in adults and larvae, as it is used for modeling epilepsy disorders by genetic manipulation [15–17]. In 2005, Baraban and colleagues used the pro-convulsant agent pentylenetetrazol (PTZ) to induce seizures in zebrafish larvae [16]. They observed a specific seizure behavior, up-regulation of the *c-fos* gene in the brain, and electrographic discharges in the optic tectum after exposing the larvae to 15 mM PTZ at seven days post-fertilization. Moreover, these responses were attenuated by common anti-seizure medications. Overall, these findings suggest that zebrafish exhibit many similarities to traditional models such as rodents.

Historically, natural substances from plants and animals are a source of numerous medicinal preparations, widely employed for preventive and therapeutic care [18,19]. This fact is corroborated by the Food and Drug Administration (FDA), which reports that between 1981 and 2019, 34% of drugs were based on substances from natural products [20].

Considering the advantages of zebrafish for drug screening based on phenotype and the need to find new treatments for controlling seizures, we aimed to evaluate the effect of different natural compounds on seizure suppression using the PTZ-zebrafish model.

## Materials and Methods

### Zebrafish maintenance and embryo acquisition

Wild-type adult zebrafish were obtained from the Laboratory of Zebrafish and Husbandry at the School of Medical Sciences, Unicamp. The animals were housed in 30 – 50 liter tanks, accommodating two animals per liter of water. The tanks were maintained under controlled physicochemical conditions of temperature (26±2°C), pH (7-7.5), levels of ammonia (< 0.1 ppm), nitrite (< 0.2 ppm) and dissolved oxygen (4-11 ppm). A photoperiod cycle of 14 h of light and 10 h of darkness was maintained. Adult fish were fed three times a day with commercial flake food (Tetramin, Tetra, Blacksburg, VA, USA) and once a day with brine shrimp and paramecium. Embryos were collected following natural spawning and nurtured in Petri dishes containing water from the aquariums maintaining consistent temperature and photoperiod with the adults. From the 5th day onwards, the larvae were fed with paramecium. Ethical approval for all experimental protocols was obtained from the Ethics Committee for Animal Research of the State University of Campinas (CEUA 4895-1/2019 and CEUA 5757-1/2021).

### Compounds

#### Tripeptide (p-BTX-I)

The compound sequence (Glu-Val-Trp) was obtained from AminoTech – Research and Development (São Paulo, Brazil). Subsequently, it was diluted in Milli-Q water (Merck KGaA, Darmstadt, Germany) and then aliquoted. These aliquots were stored at -20°C. The employed concentrations (as listed in Table 1) were determined based on research conducted in cell models [21,22].

**Table 1.** Compounds and their concentrations were analyzed to determine their effect on the suppression of seizures induced by the proconvulsant agent pentylenetetrazole.

| Compound | Concentration |
| --- | --- |
| Tripeptide (p-BTX-I) | 25, 10 and 5 µg/mL |
| CX2 | 0.5, 0.1 and 0.05 µM |

#### CX2

The CX2 (ARG-Cx43p) sequence corresponds to the C-terminal domain of the zebrafish Cx43 gene (PCT/EP2020/071242). CX2 was assembled following standard Fmoc/tBu solid-phase MW-assisted apeptide synthesis protocols [23,24], purified by reverse-phase HPLC, and their identity confirmed by HPLC-MS(ESI). The concentrations used (as listed in Table 1) were determined through studies involving zebrafish models for regeneration and senescence. Like Tripeptide, CX2 was diluted in Milli-Q water, aliquoted, and stored at -20°C.

### Compounds pretreatment

For all compounds, zebrafish larvae at 6 days post-fertilization (dpf) were randomly placed into Petri dishes, each one containing 25 larvae. The larvae were incubated in the respective solutions and concentrations for 24h (as specified in Table 1) prior to seizure induction. The temperature and photoperiod conditions were consistent with those described for the adult zebrafish. The study groups consisted of the Control Group (CG), PTZ Group (PTZ), Treatment 1, Treatment 2, and Treatment 3. Treatments 1 to 3 employed the previously determined concentrations mentioned above.

### Seizure induction by pentylenetetrazol

Seizure induction followed the pretreatment of the compounds. Larvae with 7 dpf were carefully transferred to a 96-well plate filled with 100 μL of aquarium water, with one larva per well. Subsequently, 100 μL of 30 mM of the proconvulsant agent pentylenetetrazol (PTZ) (Sigma-Aldrich, St. Louis, MO, USA) was added to each well, resulting in a final concentration of 15 mM of PTZ. The larvae were exposed to PTZ for 20 minutes. The control group underwent the same procedure but in PTZ-free water.

### Latency and number of seizures

The latency and number of seizures were monitored through visual observation, following the method described by Barbalho et al., 2016a [25]. Specifically, latency was defined as the period between the initiation of PTZ exposure and the larva reaching stage 3 of seizure-like activity, according to the criteria established by Baraban et al., 2005 [16]. Larvae were evaluated for 10 minutes of PTZ exposure, and a complete seizure was considered when they reached stage 3.

### RNA extraction

Immediately after PTZ exposure, fish were cryo-anesthetized and their heads were cut, collected, and then incubated in TRIzol® (Invitrogen, Carlsbad, CA, USA). The biological material was lysed using the TissueLyzer equipment (QIAGEN, GmbH, Germany) at 25 beats per second (BPS) for 2 minutes. Subsequently, the heads were stored at -80 °C until further processing. Each group consisted of five samples (n = 5); however, each sample was composed by pooling five larval heads, to obtain sufficient biological material for RNA extraction. Total RNA extraction was performed using TRIzol® according to the manufacturer’s protocol. The concentration and quality were determined using the Epoch™ spectrophotometer (BioTek, Winooski, VT, USA) and gel electrophoresis.

### RT-qPCR

The synthesis of cDNA was carried out using the High-Capacity cDNA Reverse Transcription kit (Applied Biosystems™, Califórnia, USA) following the manufacturer’s instructions. Quantitative PCR (qPCR) was performed using the SYBR ® Green Master Mix reagent (Bio-Rad) using the ABI 7500 system (Applied Biosystems, Foster City, CA, USA). Target genes (as listed in Table 2) were designed using the Primer-BLAST online tool (https://www.ncbi.nlm.nih.gov/tools/primer-blast/) from NCBI (National Center for Biotechnology Information) zebrafish database. Runs were carried out in triplicate using *eef1a1l1* as a housekeeping gene [26] to normalize the genes from the inflammation pathway (*il1b, tnfa, cox1, cox2a* and *il6*), the apoptosis pathway (*casp3a, casp9, baxa* and *bcl2a*), and the oxidative stress pathway (*nox1, sod* and *cat*) were analyzed. Data were evaluated using 7500 software v2.03 (Applied Biosystems). Relative gene expression analysis was calculated using the Livak and Schmittgen equation RQ = 2-ΔΔCT [27].

**Table 2:**
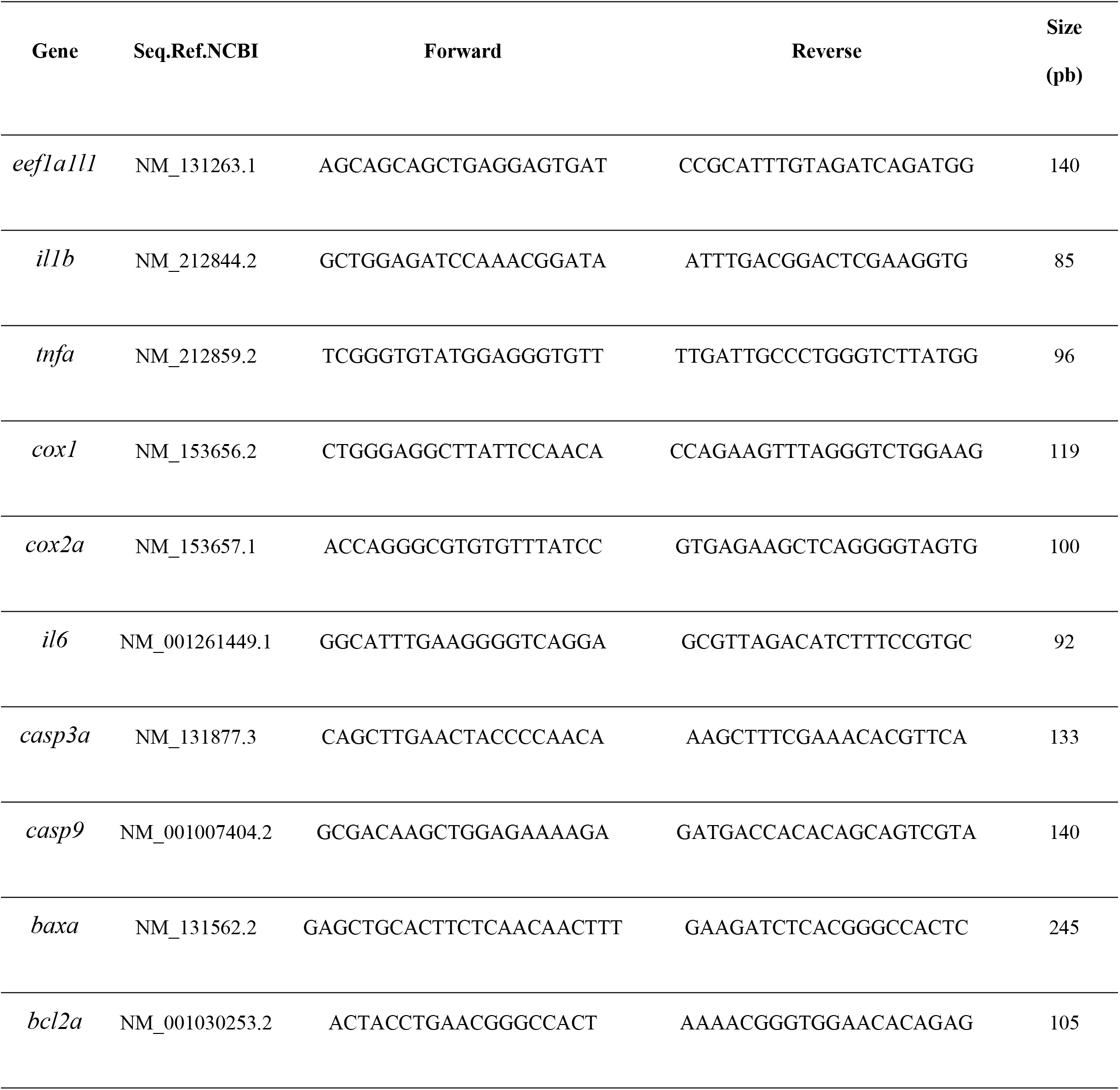
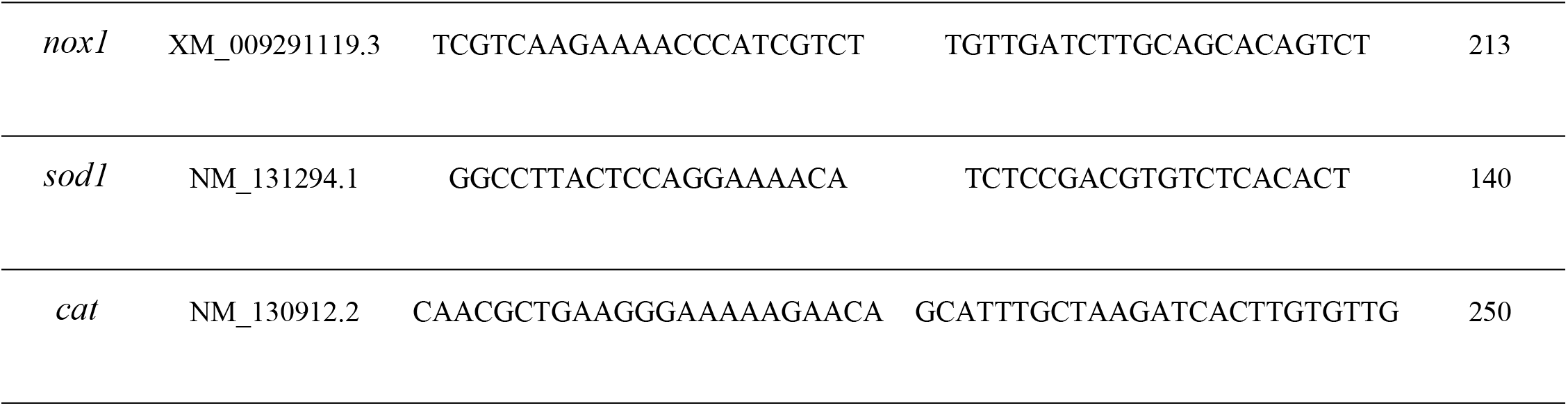
Primer sequences designed using the Primer-Blast online tool.

The three pathways (inflammation, apoptosis, and oxidative stress) are all closely linked to the underlying mechanisms of seizure generation and progression, which is why we selected representative genes from these pathways for analysis.

### Statistical analysis

Statistical analysis was performed using GraphPad Prism version 5.0 (GraphPad Software, San Diego, CA, USA). One-way ANOVA was performed, followed by the Bonferroni method for multiple comparisons. Differences were considered significant if p<0.05. Results are presented as mean ± SEM.

### Bio-distribution

Bio-distribution analysis was conducted on Tripeptide and CX2, labeled with tetramethylrhodamine-TMR and carboxytetramethylrhodamine – TAMRA, respectively.

Larvae at 6 dpf were transferred to a 96-well plate, with one larva per well, containing 100 μL of aquarium water. Subsequently, 100 μL of the Tripeptide compound at 20 μg/mL or CX2 at 1 μM was added according to the respective group, totalizing 200 ml of solution at final concentration of 10 μg/ml for Tripeptide and 0.5 μM for CX2. Exposure times for both compound groups were 1, 6, 18, and 24 hours (n=3 each time). The control group underwent similar manipulation but in water only. The incubation temperature during the experiments was maintained at 26±2°C. upon completion of exposure, larvae were rapidly and carefully transferred to a Becker containing 0.02% tricaine (Tricaine, Sigma) for approximately 2 minutes. Subsequently, the larvae were transferred to a Petri dish containing 1% agarose as a base, along with a drop of water. Larvae were examined and photographed using the Multizoon AZ100 microscope (Nikon, Tokyo, Japan) at 2X magnification. Two photographs of each larva were taken, one in black and white, and the other capturing only the fluorescence emission. Imaging merging was performed using the NIS-Elements software (Nikon, Tokyo, Japan). The experiments and protocols were approved by the animal care and use committee of the Universidad de Santiago de Compostela and the standard protocols of Spain (CEEA-LU-003 and Directive 2012-63-EU).

## Results

### Behavioral assay

We analyzed the effects of tripeptide and CX2 on seizure behavior. We found an increase in latency only for animals pretreated with the CX2 at 0.1 μM, (p≤0.05) as well as a significant decrease in the number of seizures (p≤0.001) when compared to the PTZ group (Fig 1B). No statistical significance was found for the tripeptide (Fig 1A).

**Fig 1:**
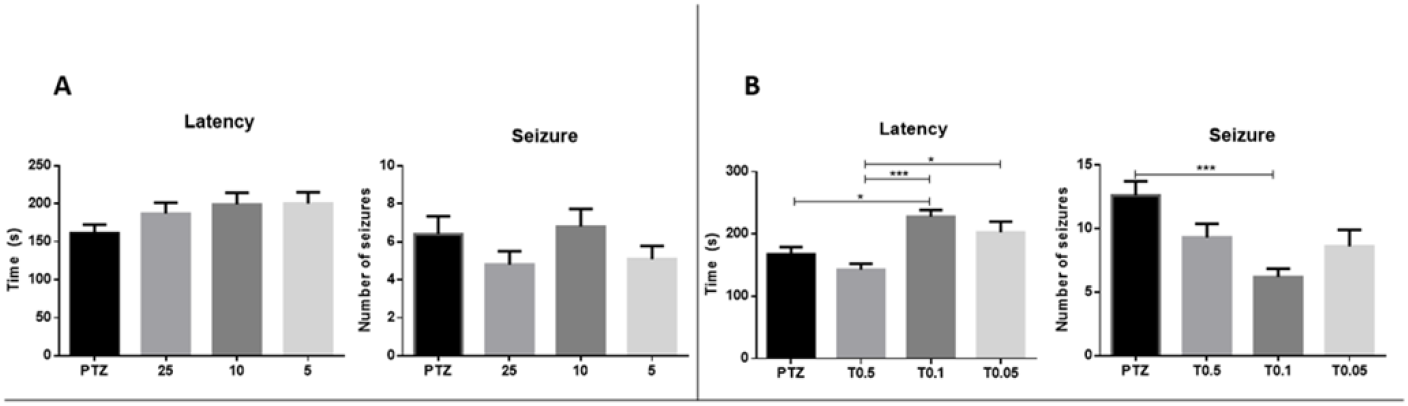
The effect of compound treatment prior to pentylenetetrazol (PTZ)-induced seizures on the number of seizure-like behaviors and latency was investigated. The number of seizures and latency were determined through visual inspection during a 10-minute exposure to PTZ (15 mM). (A) Animals exposed to the tripeptide for 24 hours: Pentylenetetrazol (PTZ) group; treatment group 25 μg/mL (T25); 10 μg/ml treatment group (T10); treatment group 5 μg/ml (T5). (B) Animals exposed to CX2 for 24 hours: Pentylenetetrazol (PTZ) group; treatment group 0.5 μM (T0.5); treatment group 0.1 μM (T0.1); treatment group 0.05 μM (T0.01). Data are presented as mean ± SEM. Statistical analyses were performed using one-way ANOVA, followed by the Bonferroni method for multiple comparisons. Differences were considered significant if p < 0.05. An asterisk (*) indicates that p≤0.05; three asterisks (***), p≤0.001.

### Molecular assay

Regarding the tripeptide, our results indicated downregulation of the *il1b* and *casp9* genes (Fig 2A and 2B), with significant effects at low concentrations (10 and 5 μg/mL). Additionally, we observed that the expression of the *casp3a, baxa*, and *bcl2a* genes was most significantly reduced at a concentration of 25 μg/mL (p<0.05), compared to the other concentrations. Similary, at a concentration of 10 μg/mL, the *casp3a, baxa*, and *bcl2a* genes exhibited notably high expression levels, roughly equivalent to or slightly higher than those of the PTZ group. However, no statistical difference was detected.

**Fig 2:**
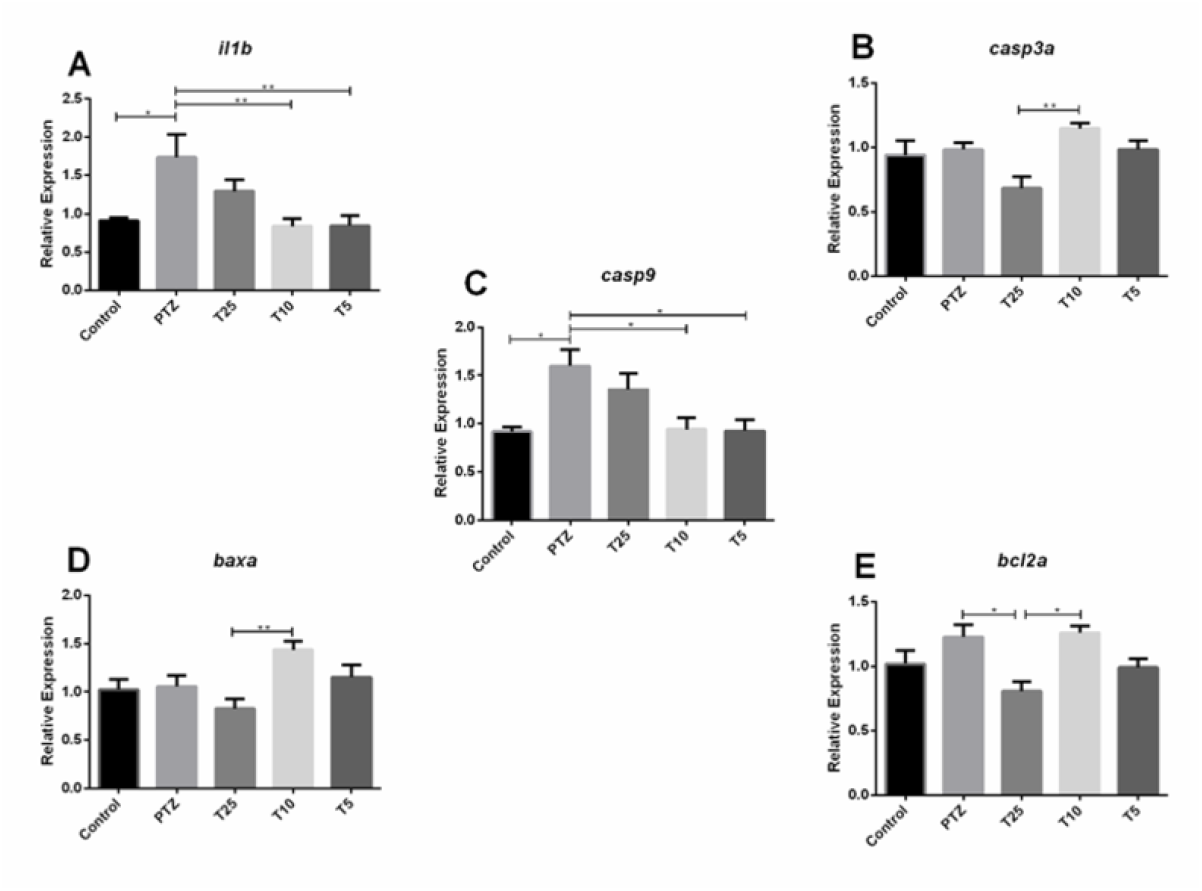
Expression of *il1b, casp3a, casp9, baxa* and *bcl2a* genes in zebrafish brain after pentylenetetrazol (PTZ)-induced seizures. Each treatment group was initially exposed to the tripeptide for 24 h and subsequently to 15 mM PTZ for 20 min, and the control and PTZ groups were handled identically, but with exposure to water (n = 5 per group). Data are presented as mean ± SEM. Statistical analyses were performed with the one-way ANOVA, followed by the Bonferroni method for multiple comparisons. Differences were considered significant if p<0.05. An asterisk (*) indicates that p≤0.05; two asterisks (**), p≤0.01. Control group (Control); pentylenetetrazol (PTZ) group; treatment group 25 μg/mL (T25); 10 μg/ml treatment group (T10); treatment group 5 μg/mL (T5).

In the case of the CX2 compounds, we found down-regulating of the *il1b, cox1, il16, and tnfa* genes for most of the analyzed concentrations (Fig 3) compared to the PTZ group. The most significant down-regulation was observed at a concentration of 0.1 μM. Conversely, the *cox2a* gene displayed increased expression across all the three concentrations tested, with the most significant increase occurring at a concentration of 0.05 μM (p≤0.001).

**Fig 3:**
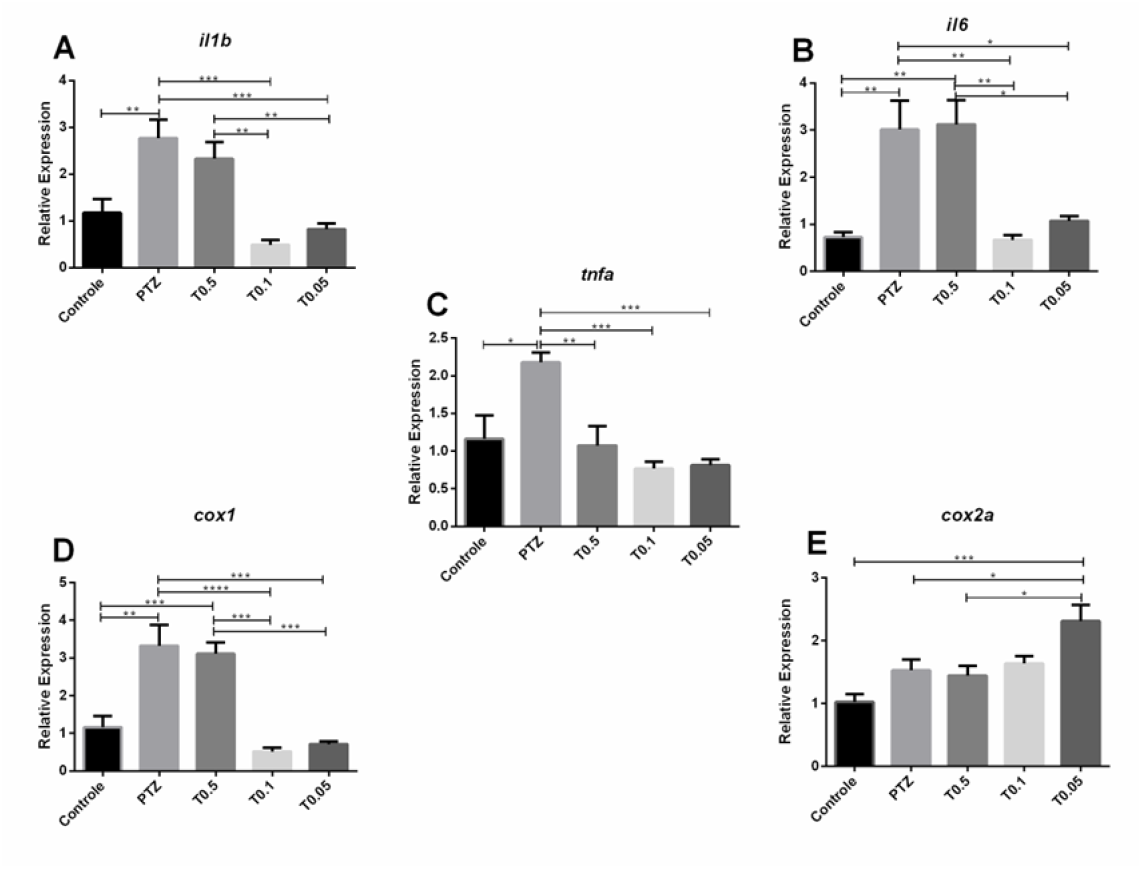
Expression of *il1b, il6, tnfa, cox1* and *cox2a* genes in zebrafish brain after pentylenetetrazol (PTZ)-induced seizures. Each treatment group was initially exposed to the CX2 for 24 h and subsequently to 15 mM PTZ for 20 min, and the control and PTZ groups were treated identically but with water exposure (n = 5 per group). Data are presented as mean ± SEM. Statistical analyses were performed with the one-way ANOVA, followed by the Bonferroni method for multiple comparisons. Differences were considered significant if p<0.05. An asterisk (*) indicates that p≤0.05; two asterisks (**), p≤0.01; three asterisks (***), p≤0.001; four asterisks (****), p≤0.0001. Control group (Control); pentylenetetrazol (PTZ) group; treatment group 0.5 μM (T0.5); treatment group 0.1 μM (T0.1); treatment group 0.05 μM (T0.01).

### Bio-distribution

By tagging both the Tripeptide and CX2 with fluorescence, we tracked their biodistribution at four time points, 1h, 6h, 18h and 24h. The CX2 compound exhibited robust detection in the brain after 1h and persisted at 6 hours (Fig 4). Additionally, the compound was detected throughout the body, indicating successful absorption by the zebrafish. In contrast, the tripeptide compound did not reach the brain, and it was predominantly retained in the zebrafish intestine (S1 Fig).

**Figure 4:**
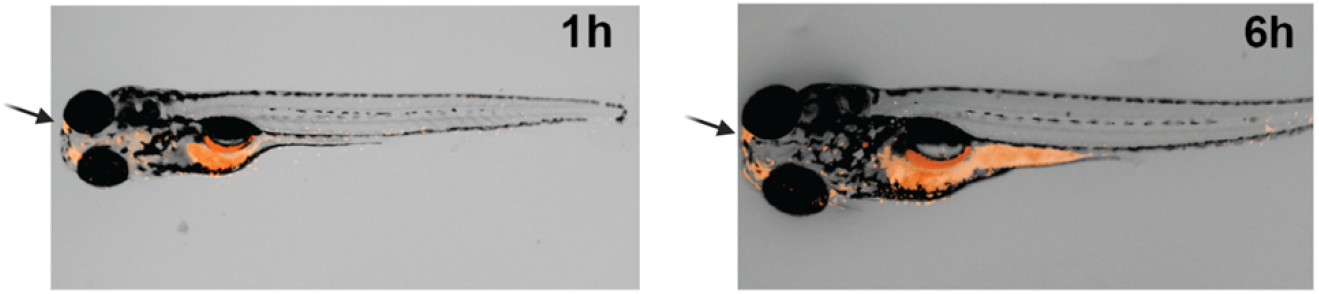
Time course of CX2 (8ARG-Cx43p, corresponding to the C-terminal domain sequence of Cx43) efficiently crossing the blood-brain barrier (BBB) in a 7-day-old zebrafish larva. The carboxytetramethylrhodamine-TAMRA labeled peptide exhibited fluorescence microscopy detection in the brain and throughout the body (indicated by red color) at 1 hour and 6 hours post-treatment.

## Discussion

Despite the availability of pharmacological treatments, uncontrolled seizures remain a concern, requiring further investigations for new seizure-suppression approaches [28]. Seizures occur due to a complex process involving multiple factors and affecting various cellular pathways [29]. This study focused on targeting inflammation, oxidative stress, and cell death pathways for seizure modulation [30–33]. To achieve our goal, we used tripeptide (p-BTX-I) to address inflammation and cell death, and the CX2 for its anti-inflammatory action[21,22].

Zebrafish offer many advantages in drug screening and phenotype assessment. The zebrafish PTZ-seizure model mimics human seizure behavior and electrographic patterns and is responsive to anti-seizure drugs. The scalability and rapid results of the zebrafish make them an advantageous model for discovering new anti-seizure compounds [8,9,16,34,35].

The process of inflammation involves the recruitment of many inflammatory mediators such as interleukins (ILs), interferons (IFNs), tumor necrosis factors (TNFs), and growth factors. *IL1B, TNF*-α, and *IL6* genes are among the most studied inflammatory cytokines in the CNS [32]. After a seizure, genes such as *IL1B, IL2, IL6, TNF*-α, and *VEGF*, which are normally present in low concentrations, increase rapidly, leading to damaging changes in synapses and heightened neuronal excitability [33,36].

Prostaglandins/*COX-2* are also produced, potentially disrupting the blood-brain barrier and significantly contributing to seizure onset and recurrence [37].

Among the compounds analyzed, CX2 at a concentration of 0.1 μM showed the most significant results in seizure suppression, reducing seizure frequency and increasing latency (Fig 1B). Treatment with CX2 prior to PTZ-induced seizures down-regulated the *il1b, cox1, tnfa, and il16* genes, indicating its role in regulating inflammation during seizures [33]. In contrast, the *cox2a* gene showed up-regulation compared to the PTZ group (Fig 3E). *COX2*, which is responsible for prostaglandin production [38], increases during seizures [39]. While *COX2* inhibitors have been explored as therapeutic agents; controversies exist due to types of inhibitors and timing of administration [39,40]

Studies in rats and zebrafish suggest that *cox1* may play a more crucial role in inhibiting seizure [41–43]. Barbalho et al. found that inhibiting *cox-1* had a positive impact on seizure in zebrafish larvae, while *cox-2* inhibition did not affect seizures [25].

Our results align with these findings. CX2 decreased the levels of *cox1* transcript (Fig 3D) and reduced the occurrence of seizures by increasing latency and decreasing their frequency (Fig 1B). The compound was quickly absorbed and distributed, as it was detected in the brain at the earliest time point examined (Fig 4), and also in the optic tectum, a region associated with significant neuronal activity during seizures [44].

As for the Tripeptide compound (Fig 2), we observed various gene responses, particularly the down-regulation of genes related to inflammation. Neuroinflammation, triggered by factors such as tissue damage, infection, stress, and seizures, has been associated with epilepsy [28,45]. Pathways connecting epilepsy to neuroinflammation have been identified, with animal models of seizure induction showing increased mRNA levels of genes involved in inflammatory cascades [46–48]. Thus, efforts have focused on low molecular weight compounds acting on these pathways. Tripeptide (10 μg/mL) downregulated *il1b* gene and *casp9* gene indicating its positive effect on these pathways.

Regarding the bio-distribution of the Tripeptide compound, it was primarily detected in the zebrafish larvae intestine (S1 Fig). This could be attributed to the fluorophore, tetramethylrhodamine-TMR, which along with the compound, may interfere with absorption or crossing the blood-brain barrier for CNS delivery.

## Conclusion

Our study provides evidence that Tripeptide and CX2 – Cx43 peptide have the potential to modulate gene expression and reduce epileptic seizures in zebrafish. Tripeptide increased the expression of genes associated with antioxidant functions while decreasing the expression of inflammatory genes, with no observed behavioral changes. The compound CX2 peptide demonstrated high effectiveness in reducing the expression of inflammatory genes, lowering the number of seizures, and increasing latency time. CX2 was widely distributed in the zebrafish brain and the body. These compounds could represent a promising avenue for further research in the development of novel anti-seizure medications. Nevertheless, further studies are needed to assess their effects in other animal models and determine their efficacy in humans.

## Supporting information

**S1 Fig.**
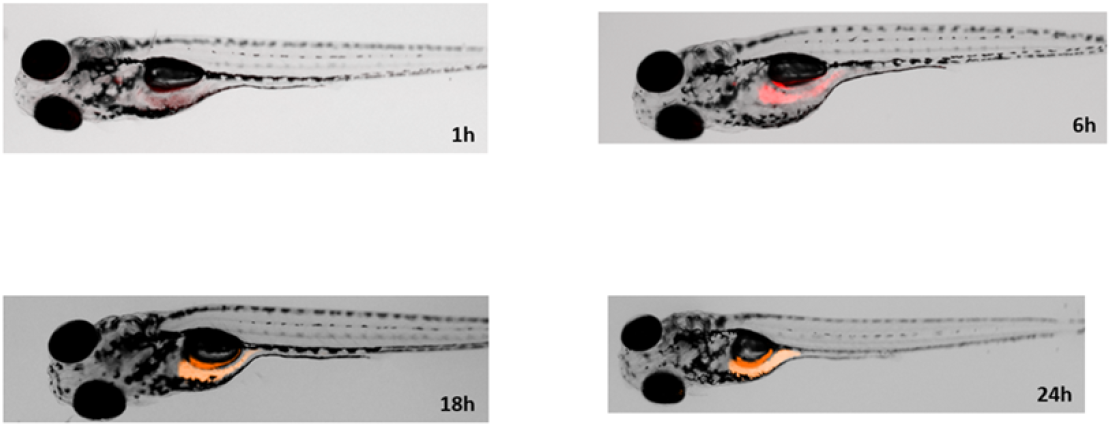
Time course of the tripeptide in a 7-days post fertilization zebrafish larva. The tetramethylrhodamine-TMR labeled peptide was not detected in the brain through fluorescence microscopy.

